# A Double Dissociation between Savings and Long-Term Memory in Motor Learning

**DOI:** 10.1101/2022.08.18.504373

**Authors:** Alkis M. Hadjiosif, J. Ryan Morehead, Maurice A. Smith

## Abstract

Both declarative and procedural memories are easier to reacquire than learn from scratch. This advantage, known as savings, has been widely assumed to result from the reemergence of stable long-term memories. In fact, the presence of savings has often been used as a marker for whether a memory had been consolidated. However, recent findings have demonstrated that motor learning rates can be systematically controlled, providing a mechanistic alternative to the reemergence of a stable long-term memory, and recent work has reported conflicting results about whether implicit contributions to savings in motor learning are present, absent, or inverted, suggesting a limited understanding of the underlying mechanisms. In order to elucidate the mechanism responsible for savings in motor learning, we investigate the relationship between savings and long-term memory by determining how they depend on different components of motor learning. To accomplish this, we experimentally dissect motor adaptation based on short-term (1-minute) temporal persistence. Surprisingly, we find that a temporally-volatile component of implicit learning leads to savings whereas temporally-persistent learning does not, but that temporally-persistent learning leads to long-term memory at 24 hours whereas temporally-volatile learning does not. Moreover, we find that temporally-persistent implicit learning not only fails to contribute to savings, but that it produces an anti-savings which acts to reduce the net savings, and we show that the balance between temporally-volatile and temporally-persistent components can explain seemingly inconsistent reports about implicit savings. The clear double dissociation between the mechanisms for savings and long-term memory formation challenges widespread assumptions about the connection between savings and memory consolidation, and provides new insight into the mechanisms for motor learning.

## Introduction

Memories, both declarative and procedural, are easier to reacquire than to learn from scratch. This advantage, known as savings, was first appreciated in Hermann Ebbinghaus’s seminal work[1], in which he observed that relearning a forgotten list of words is faster than learning a novel list. Savings has since been demonstrated in a plethora of different paradigms, including cognitive tasks in humans[2,3], operant conditioning in animals[4–6], and motor tasks in humans such as saccade adaptation[7], force-field adaptation[8–11], visuomotor adaptation[12–20] and gait adaptation[21,22].

Previous research generally maintained that savings results from the recall of a previously consolidated memory. In fact, the presence or absence of savings itself has often been taken as a litmus test for whether a previously trained memory has been consolidated[14,23,24]. In line with this idea, savings has been further suggested to result from (1) the unmasking of a slower-learning, strong-retention process in a multi-rate learning model[10]; (2) context- or relevance-based switching between such multiple slow processes, each specific to a different memory[25–28]; or (3) reverting to the memory of a previously learned motor plan that was reinforced by success or mere repetition[16,29]. All of these proposed mechanisms focus on savings as the manifestation of a latent, stable, consolidated motor memory that is robust to both interference and the passage of time.

In line with this idea, a recent but influential view has proposed that savings in motor adaptation specifically results from the recall of an explicit strategy [18,30,31], whereas the implicit component of visuomotor learning does not contribute to savings[31]. These studies provided clear evidence for explicit savings, but the paradigms they used elicited little implicit adaptation, limiting the power to assess implicit savings. More recent studies, which elicited greater implicit adaptation, have led to disparate findings, concluding that implicit adaptation is either faster during relearning[19,20] or, actually, slower[32]. The Avraham and Albert studies are of particular interest, as they were both designed to *isolate* implicit adaptation (albeit using different paradigms) and yet reached the opposite conclusion.

What could explain this apparent discrepancy? A possibility is that implicit adaptation may not be monolithic. It may consist of distinct components that are reacquired at different rates and are differentially elicited in these different paradigms. In fact, an intriguing alternative to recall-based mechanisms is that savings arises from such changes in learning rate[11,15,22], an idea reinforced by recent work which has demonstrated that the rate at which learning occurs is systematically modulated by specific characteristics of the learning environment. These characteristics include the amount of task-relevant variability present before learning[33,34], the balance between sensory uncertainty and uncertainty about state estimation[35,36], prior exposure to perturbations characterized by a similar covariance structure between learning parameters[37,38], prior exposure to similar motor errors[39], and the trial-to-trial consistency of the learning environment[40,41]. In particular, high consistency environments, whereby perturbations tend to persist from one trial to the next and thus confer more predictability to the imposed perturbation, can strongly increase learning rates (up to 3x). This is a critical finding as far as the study of savings is concerned: The adaptation paradigms used to study savings usually consist of the same perturbation being active for a large number of trials (usually 60-120), resulting in high environmental consistency, which would in turn lead to strongly increased learning rates. This mechanism could lead to savings by enabling the faster re-acquisition of a short-term memory of learning, as opposed to the reemergence of a long-term, stable memory of this learning.

Here, we compare the mechanisms that lead to savings, and those that lead to the formation of stable, long-term motor memories. We hypothesized that dissecting motor adaptation into specific memory components based on temporal persistence could shed light into these mechanisms. We use short 1-minute time delays to dissect overall adaptation into two components: temporally-volatile adaptation, which would decay during this time delay, and temporally-persistent adaptation, which would survive the delay[42–44]. Surprisingly, we find that savings is driven not by the reemergence of temporally-persistent motor memories, but instead by faster relearning of temporally-volatile memories. We go on to find that these temporally-volatile memories responsible for savings in our paradigm represent implicit, rather than explicit, adaptation. When we measure the long-term retention of the temporally-persistent and temporally-volatile components, however, we find that it is temporally-persistent adaptation, not temporally-volatile adaptation, that leads to long-term memory. Together, these findings demonstrate an intriguing double dissociation between savings and long-term memory.

## Results

We designed a set of experiments to elucidate the mechanisms for savings and long-term memory and investigate the relationship between them. We began by investigating whether savings, the faster reacquisition of a previously learned adaptation, is driven by the re-emergence of a previously consolidated temporally-persistent memory, or by a propensity for faster acquisition of a transient, temporally-volatile memory. In particular, we created a paradigm to dissect initial adaptation, the washout of adaptation, and savings in readaptation into temporally-persistent and temporally-volatile components. We first investigated the dynamics by which temporally-persistent and temporally-volatile memories decay during a washout period following initial adaptation, as savings can arise from the incomplete washout of a component of adaptation[10,45]. This allowed us to compare the rates of unlearning for temporally-persistent and temporally-volatile adaptation during washout, and critically, to measure the initial value of both temporally-persistent and temporally-volatile memories prior to readaptation, so that savings could be accurately assessed for both. We next examined how savings depends on temporally-persistent and temporally-volatile memories by measuring savings separately for these two components of motor adaptation, allowing us to determine whether one of these memories is specifically responsible for savings.

For Experiments 1 and 2, we recruited N = 40 subjects and trained them on a 30° visuomotor rotation (VMR)[12,13,15–18,46–48] (**Fig. 1a,b**). After 80 trials of initial training, subjects were tested for savings after either a short (40-trial) washout period, which was previously reported to be sufficient for the washout of overall VMR adaptation[15], or a longer (800-trial) washout period we employed to effect a more definitive washout. We also used the data from this 800-trial washout to trace out the time-course of unlearning for both the temporally-persistent and temporally-volatile components of adaptation; this unlearning would encompass both active unlearning (i.e. relearning the baseline behavior) and natural trial-to-trial decay of adaptation[17,49–52]. Each subject experienced both types of washout duration following training (see **Fig. 1c**). In Experiment 1 (N=20), the short washout period was presented first and the long washout period second. In Experiment 2 (N=20), this order was flipped (**Fig. 1c**, for a detailed description see Materials and Methods).

**Figure 1.**
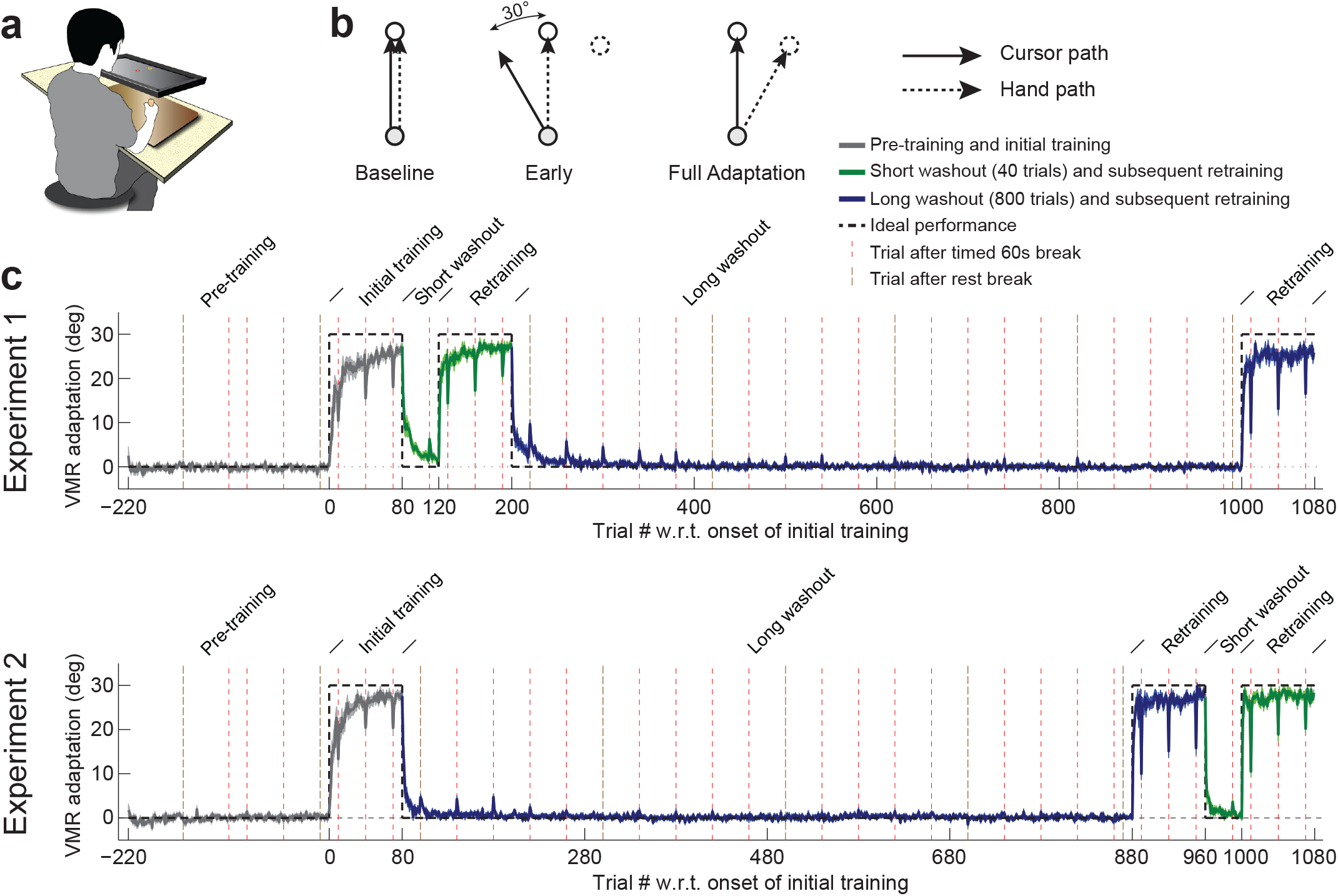
Experiment setup and training paradigm. **(a)** Experiment setup. Subjects made point-to-point reaching movements on a digitizing tablet and received continuous visual feedback on a screen mounted above it. **(b)** Visuomotor rotation training. During baseline (left), the cursor follows the hand motion, whereas during training cursor motion is skewed by 30° from the hand motion (in this example, counter-clockwise), resulting in a 30° error before adaptation (middle). If full adaptation is achieved, hand motion must completely counter the imposed rotation, corresponding to a 30° clockwise hand motion in this example (right). **(c)** Top: experiment schedule and raw data for Experiment 1. There were three phases: a baseline period followed by the initial 80-trial VMR training (average adaptation level shown in gray); a short, 40-trial washout period followed by retraining (green); and a long, 800-trial washout period followed by another 80-trial retraining session (blue). Red dashed vertical lines indicate trials conducted after 60-second breaks to isolate temporally-persistent adaptation. Brown dashed vertical lines indicate trials following rest breaks. Note that, during the washout periods, adaptation peaks on these break trials, illustrating a slower washout for temporally-persistent vs. temporally-volatile adaptation. Bottom: same but for Experiment 2, where the long, 800-trial washout period came first. Errorbars indicate SEM.

### Persistent adaptation washes out more slowly than overall adaptation

The data from the long, 800-trial washout period allowed us to carefully examine the time-course of unlearning for both the overall adaptation and for the temporally-persistent component of it. We measured the latter by occasionally inserting one-minute breaks (every 40 trials) which would allow for temporally-volatile adaptation to decay. Since the one-minute breaks we imposed amount to 3-4x the time constant for decay of temporally-volatile adaptation[42,44], 95-98% decay of volatile adaptation would be expected, effectively isolating temporally persistent adaptation.

Analysis of the washout curves revealed that overall adaptation displayed rapid unlearning; however, persistent adaptation (circles in **Fig. 2a**) was unlearned much more slowly. We found that by trials 16-25, labeled as “early washout” in **Fig. 2a**, overall adaptation had already dropped below 10% of the pre-washout asymptotic adaptation level, whereas about 40% of pre-washout persistent learning remained. By trials 51-150, labeled as “mid washout”, overall adaptation had dropped below 3%, whereas about 20% of persistent adaptation still remained (**Fig. 2a**, see inset). Correspondingly, we found the retention of persistent learning to be significantly greater than overall learning in both early washout (t(23) =4.8, p = 6.9 × 10^-5^ and mid washout periods (t(39) = 8.5, p=1.9 × 10^-10^). To quantify the rate of unlearning during washout for both overall and persistent adaptation, we fit single exponential decay functions to the washout data (see Materials and Methods). This revealed the time constants for unlearning to be 6-fold slower for temporally-persistent adaptation than for overall adaptation (median time constant estimated using bootstrap: 106.0 trials, IQR [92.6 to 121.9] vs. 17.4 trials, IQR [15.1 to 20.1], p<10^-4^, **Fig. 2a**), in line with the higher retention we observed in the early- and mid-washout data.

**Figure 2.**
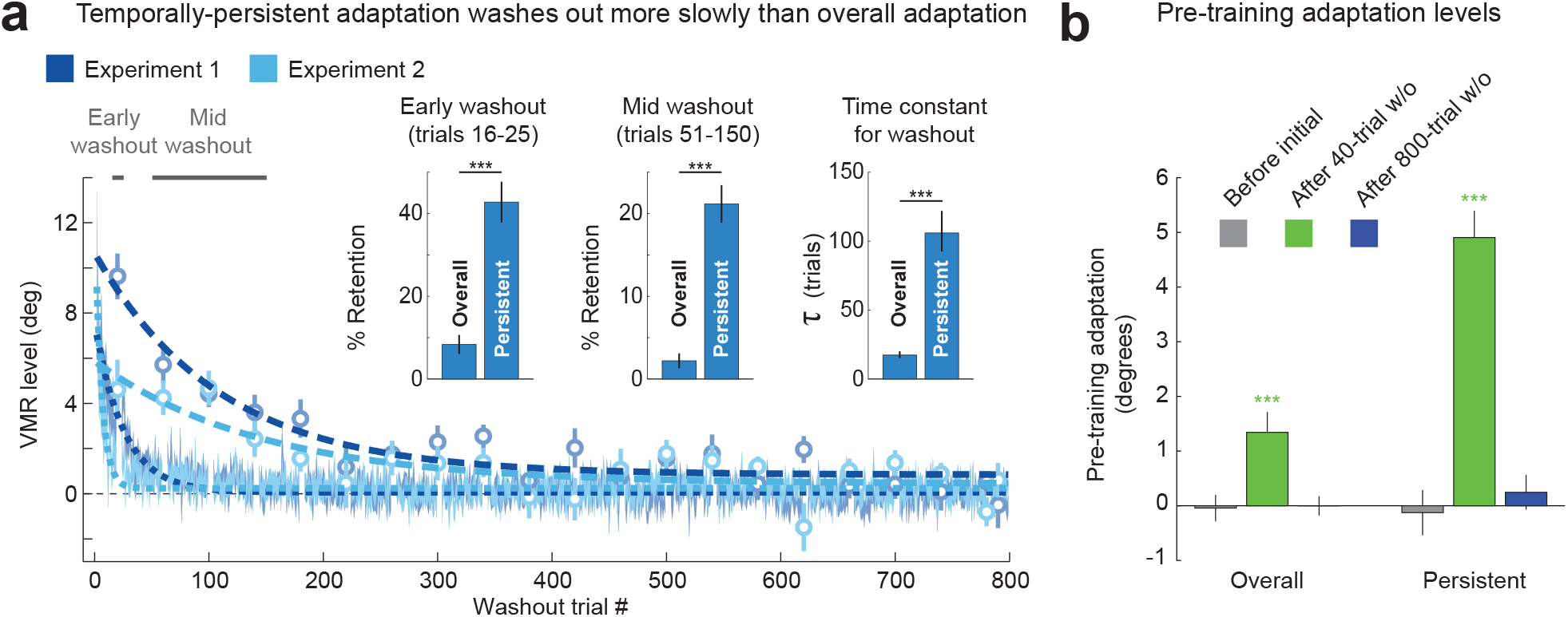
Temporally-persistent adaptation washes out more slowly than overall adaptation. **(a)** Washout curves for the overall adaptation (shading indicates mean±SEM) and temporally-persistent adaptation (circles) for both Experiment 1 (blue) and Experiment 2 (light blue), illustrating the contrast between rapid washout for overall adaptation and slower washout for temporally-persistent adaptation. The thick dashed or dotted lines indicate exponential fits. Inset: Three different measures of washout for overall vs. persistent adaptation. Retention expressed as a percentage of asymptote adaptation after 16-25 trials (left) or 51-150 trials (center) indicates slower washout for temporally-persistent compared to overall adaptation. Time constants for the washout curves (right) also show slower washout for temporally-persistent adaptation. *p<0.05, **p<0.01, ***p<0.001. **(b)** Residual adaptation before initial training (gray, left bar in each cluster), at the end of the 40-trial washout period (green, middle bar) and at the end of the 800-trial washout period (blue, right bar). The data show that the 40-trial washout period leaves a significant amount of temporally-persistent adaptation, and a smaller but also significant amount of overall adaptation. Consequently, retraining after only 40 washout trials starts from a nonzero baseline. *p<0.05, **p<0.01, ***p<0.001.

As a minor point, we also noticed that unlearning during the 800-trial washout period was somewhat slower in Experiment 1 than Experiment 2 (Experiment 1: time constants of 108.9 trials, IQR [89.1 to 132.8] for temporally persistent and 25.1 trials, IQR [22.2 to 29.2] for overall, compared to Experiment 2: time constants of 142.2 trials, IQR [114.8 to 186.6] for temporally-persistent and 5.8 trials, IQR [4.6 to 7.1] for overall),. This might reflect the difference in the amount of training between the two conditions, as two training blocks (160 trials in total) preceded this washout period in Experiment 1, whereas only a single training block (80 trials) preceded this washout in Experiment 2, due to the condition balancing (see **Fig. 1**). In summary, we found that temporally-persistent adaptation is unlearned at a considerably slower rate than overall adaptation.

### Residual adaptation prior to the onset of retraining

One consequence of the slower unlearning of persistent compared to overall adaptation is that, while the washout of overall adaptation can appear complete after a short 40-100 trial per direction washout period [15,16,18,23], substantial temporally-persistent adaptation can nevertheless remain. This suggests that longer washout periods may be required to examine savings independent of the effect of residual temporally-persistent adaptation, and that measuring this residual adaptation prior to relearning may facilitate a better understanding of relearning behavior.

When we measured the residual overall and persistent adaptation before the onset of retraining, we found significant levels of both overall and temporally-persistent adaptation for the 40-trial washout but no significant residuals of the previous adaptation following the 800-trial washout period. In particular, we found small but significant residuals for overall adaptation at the end of the 40-trial washout periods, around 5% of pre-washout levels (1.34±0.37°, t(39) = 3.6, p = 0.00087 for Experiments 1 and 2 combined, with positive values indicating adaptation in the direction of the previously imposed VMR, **Fig. 2b**). Note that these residuals were related to previous adaptation rather than a movement direction bias because both Experiments 1 and 2 were balanced, with 10 participants trained with clockwise, and 10 with counter-clockwise VMRs for each experiment. The residuals were even larger for temporally-persistent adaptation, in line with the substantially slower unlearning of temporally-persistent adaptation compared to overall adaptation we observed. In particular, we found that the residual persistent adaptation before the end of the 40-trial washout was around 25% of pre-washout persistent adaptation (4.91±0.49° for Experiments 1 and 2 combined, t(39)=10.0, p = 2.7 × 10^-12^). In contrast, the 800-trial washout period was sufficient to bring both overall adaptation and temporally-persistent adaptation back to baseline, with measured residuals of only 0-2% of pre-washout levels on average. These residuals were not consistently in the direction of the pre-washout adaptation and were not statistically significant. Overall adaptation at the end of the 800-trial washout was −0.00±0.18°, whereas temporally-persistent adaptation was 0.25±0.32°, as shown in **Fig. 2b**.

These results show that a prolonged washout period is required to eliminate residual temporally-persistent adaptation. As washout periods in previous experimental work on savings[15,16,18,23,48] are typically much shorter than the 800-trial washout period we examined, it is likely that the savings observed in these studies is, at least in part, driven by interactions between different components of adaptation that were not fully washed out prior to retraining – *apparent* savings – as suggested in Smith et al., 2006[10]. In order to examine faster relearning that is not contaminated by such interactions – that is, examine *true* savings – one should ideally eliminate residual levels of overall, temporally-persistent, and temporally-volatile adaptation, or, at least, take these residual levels into account.

### Savings is present even when previous adaptation is completely washed out

To investigate savings for overall and temporally-persistent adaptation, we compared the learning curves for retraining and initial training, shown in **Fig. 3a,b** (gray: initial training; green: retraining after 40 washout trials; blue: retraining after 800 washout trials). We found that the adaptation levels achieved in the early adaptation period (trials 8-12 after perturbation onset, excluding trial 10 which was after a time delay), when learning was most rapid, were noticeably higher for relearning (24.6±0.7° overall; 24.9±0.6° and 24.4±1.0° after the 40-trial and 800-trial washout periods, green and blue lines, respectively, with data combined across Experiments 1 and 2 in all cases) compared to initial training (18.8±1.2°, gray). Because pre-training adaptation levels were not identical across conditions as shown in **Fig. 2b** (1.34±0.37°, −0.00±0.18°, and −0.04±0.24°, after short washout, long washout, and before initial adaptation), we normalized data to quantitatively compare learning and relearning curves independent of the effect of this residual pre-training adaptation. Specifically, we subtracted the pre-training adaptation separately for overall and temporally-persistent adaptation, and normalized each baseline-subtracted learning curve by the distance between baseline and the ideal adaptation level (Equation 2, see Materials and Methods). These normalized data, plotted in **Fig. 3c,d** express adaptation levels as a percentage of that required for full adaptation. In particular, normalized early adaptation (trial 10 after perturbation onset) was faster compared to initial adaptation (initial adaptation: 62.8±4.1% vs. relearning: 81.7±2.5%; 40-trial and 800-trial washout data separately: 82.1±2.1% and 81.4±3.2%, respectively). We defined savings simply as the difference between these normalized adaptation data for the retraining vs. the initial learning conditions. The top panel in **Fig. 3e,f** shows an estimate of this savings measure in the overall adaptation. We find statistically significant savings for early adaptation (trial 10; savings of 18.9±3.3% of the ideal adaptation, t(39)=5.8, p = 5.4 × 10^-7^ [40-trial washout data: 19.3±3.3%, t(39)=5.8, p = 4.4 × 10^-7^; 800-trial washout data: 18.6±3.6%, t(39)=5.1, p = 4.1 × 10^-6^). Inspection of the overall adaptation data in the top panels of **Fig. 3c,d** reveals that both the readaptation and initial adaptation curves asymptote near the ideal adaptation level, meaning that the room for improvement, and thus the capacity for savings, is reduced as training proceeds. In line with this observation, the savings we observed for mid (trial 40) and late (trial 70) overall adaptation were smaller than the savings observed for early (trial 10) adaptation (<10% of the ideal adaptation in all cases). The savings we observed at trials 40 and 70 were, however, statistically significant (t(39)=2.6, p = 0.0063 for trial 40 and t(39) = 3.3, p=0.0011 for trial 70), as shown in **Fig. 3e,f**.

**Figure 3.**
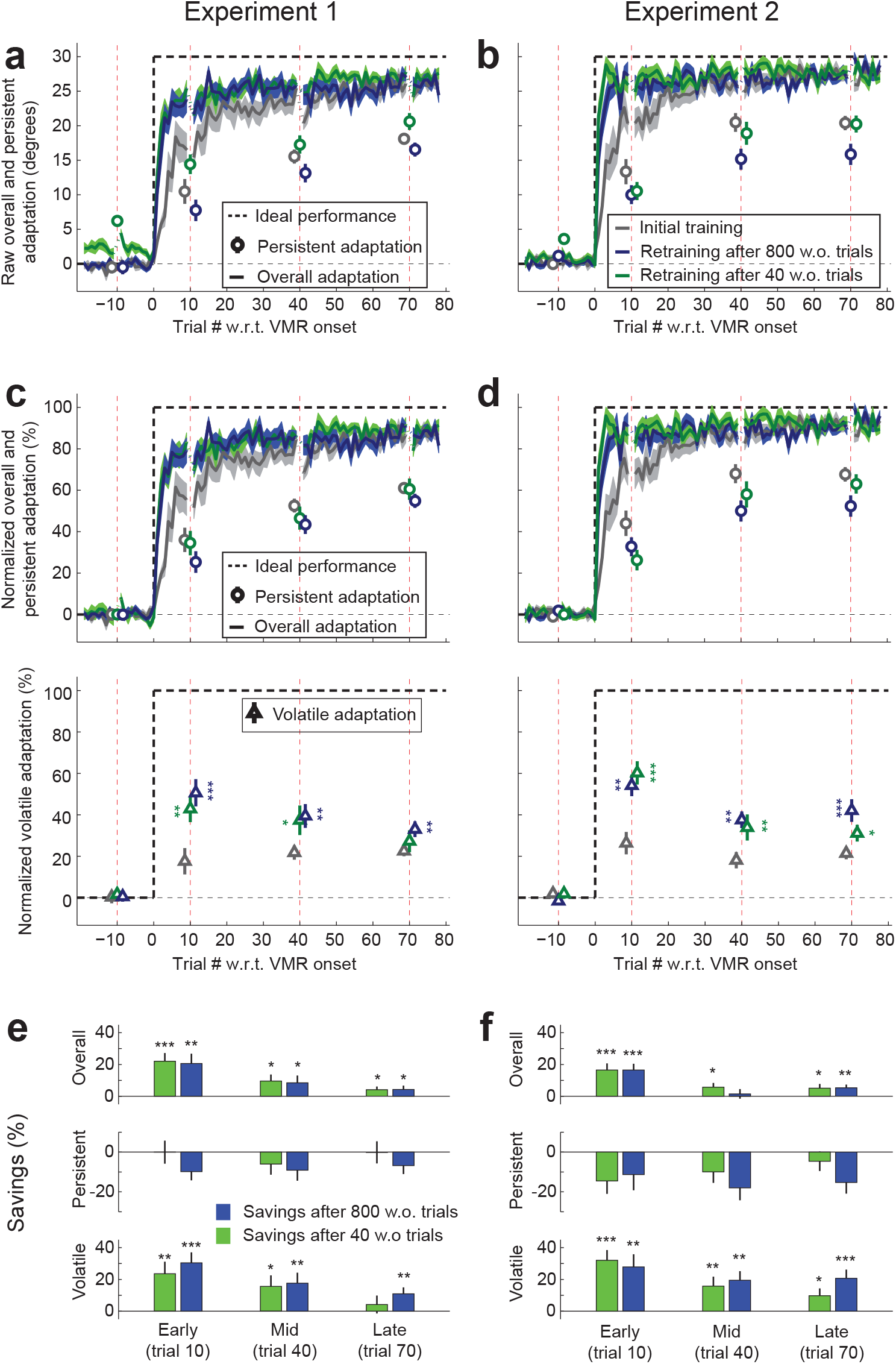
Temporally-volatile adaptation displays savings, but temporally-persistent adaptation does not. **(a,b)** Learning curves for Experiments 1 and 2, showing overall and persistent adaptation (solid lines and circles, respectively). Persistent adaptation during training or retraining was assessed after a 60-second break as shown in Figure 1. The dashed black line indicates ideal performance and errorbars indicate SEM. Overall adaptation (solid lines) is faster for retraining after both a 40-trial washout period (green) and an 800-trial washout period (blue) when compared to initial adaptation (grey). In contrast, persistent adaptation (circles) is not faster during retraining than initial training. Note that performance during the retraining period following a 40-trial washout (green) can be inflated because this washout is often incomplete for both overall and persistent learning, as indicated by nonzero pre-training levels as also shown in **2b**. Washout for both overall and persistent adaptation is complete for the 800-trial data (blue) in both Experiments 1 and 2 resulting pretraining levels that are essentially zero in all cases. **(c,d)** Normalized overall, persistent, and volatile adaptation components in Experiments 1 and Experiment 2. Upper panels show the normalized learning curves for overall and temporally-persistent adaptation. Here, the raw data shown in A were normalized by subtracting out pre-training adaptation levels so that training-related changes in performance can be directly assessed and scaling ideal performance to 100% (see Materials and Methods, Equation 2). Whereas the overall adaptation is faster for retraining after both washout periods, persistent adaptation is not. Lower panels show the normalized learning curves for temporally-volatile adaptation. Temporally-volatile readaptation is consistently higher than initial adaptation, especially for the early (trial 10) timepoint, where savings should be most pronounced and where readaptation is between 2-3 times higher. **(e,f)** Savings for overall, temporally-persistent, and temporally-volatile adaptation in Experiments 1 and 2, defined as the difference between the normalized adaptation metrics displayed in panels **c,d** for relearning vs. initial learning (see Materials and Methods, Equation 3). Positive values for savings indicate faster relearning. We find substantial and significant savings especially early during training (trial 10) for both overall and temporally-volatile adaptation, but not for temporally-persistent adaptation, suggesting that savings is driven by temporally-volatile adaptation.

### Savings does not arise from the rapid reemergence of temporally-persistent memories

Intriguingly, the clear pattern of savings we found in the learning curves for overall adaptation was not present for temporally-persistent adaptation. In only one of the four conditions in Experiments 1 and 2 (readaptation after a 40-trial washout in Experiment 1) was the persistent adaptation higher during relearning than initial adaptation, and in that condition the readaptation built upon a substantially higher pre-training level than the corresponding initial training condition (**Fig. 3a**). When pre-training levels of persistent adaptation were taken into account by normalizing learning curves, we found that relearning for temporally-persistent adaptation was slower, rather than faster, than initial learning as shown in **Fig. 3a,b**. This absence of savings is illustrated in **Fig. 3c,d**. In fact, these learning curves showed a tendency for anti-savings (slower persistent readaptation) for both washout conditions. Specifically, early (trial 10) savings were, on average −10.1±4.3% of the ideal persistent adaptation, t(38)=-2.4, p = 0.99 for savings (40-trial washout data: −7.3±4.4%, t(37)=-1.7, p = 0.95; 800-trial washout data: −10.6±4.5%, t(38)=-2.3, p = 0.99) as shown in **Fig. 3e,f**. The temporally-persistent adaptation measured 40 and 70 trials into the training period displays results similar to trial 10 adaptation, with either no savings or a tendency towards negative savings (t(39)=-3.4, p = 1.00 for trial 40 and t(38) = −2.2, p = 0.99 for trial 70).

A post-hoc analysis asked whether temporally-persistent adaptation actually displayed slower relearning. It revealed that after both washout periods combined, savings in persistent adaptation was, in fact, significantly negative (t(38) = −2.4, p=0.0232, 2-tailed paired t-test). This indicates that, for temporally-persistent adaptation, relearning was significantly slower than initial learning. Individually, This was most clear in the long 800-trial washout data (which allowed us to examine savings without any effects of residual temporally-persistent adaptation), with significantly negative savings (t(38) = −2.3, p=0.0244, 2-tailed paired t-test). Savings after the short incomplete washout was also nominally negative but, in this case, not significantly so (t(37) = −1.7, p = 0.1073, 2-tailed paired t-test). The negative savings results we observe in the 800-trial and 40-trial washout data are similar but it is likely that the 800-trial result is more reliable as the 40-trial washout data suffer from incomplete washout of the initial adaptation before relearning. Thus our data show a conspicuous absence of savings in the reacquisition of temporally-persistent adaptation in all conditions we examined, instead showing clear anti-savings despite robust savings in the reacquisition of overall adaptation.

### Savings arises from the faster acquisition of temporally-volatile memories

The absence of savings in the temporally-persistent component of adaptation suggests that the temporally-volatile component of adaptation is responsible for the savings observed in overall adaptation. We calculated savings for temporally-volatile adaptation (bottom row in **Fig. 3e,f**) based on the normalized volatile adaptation (bottom row of **Fig. 3c,d**), which was computed as the difference between the normalized overall and normalized persistent adaptation (top row of **Fig. 3c,d**). We found that volatile adaptation during early training (trial 10) was 2-3fold faster for retraining than for initial training after both the short and long washout periods in both experiments (as shown in the bottom row of **Fig. 3c,d**). Specifically, we found that trial 10 volatile readaptation was 52.4±3.8% of the ideal adaptation to the 30° VMR with data after both types of washout combined vs. 22.0±4.2% for initial adaptation, t(38)=6.2, p = 1.3 × 10^-7^ (readaptation for 40-trial washout data: 51.3±4.5%, t(37) = 5.7, p = 9.0× 10^-7^ for savings; readaptation for 800-trial washout data: 52.4±4.1%, t(38) = 5.6, p = 8.7 × 10^-7^ for savings). This indicates substantial, statistically significant savings in temporally-volatile adaptation as illustrated in the bottom row of **Fig. 3e,f** that stands in stark contrast to the absence of savings observed in temporally-persistent adaptation, suggesting that overall savings arises from the former, but not the latter.

In line with the above, it is statistically clear – both for combined data but also for 800-trial and 40-trial washout data separately – that overall savings is overwhelmingly driven by savings in volatile adaptation. In particular, when we analyzed the ratio of overall savings that is accounted for by volatile vs. persistent savings, we found that temporally-volatile savings accounted for essentially all overall savings (95% confidence intervals for the % contribution of volatile savings to overall: [110% to 219%], confidence interval estimates obtained using bootstrap, see Materials and Methods). This was similar for both the 40-trial data [95% to 192%] and the 800-trial data [112% to 235%] separately. Correspondingly, we found the contribution of temporally-persistent savings to be overwhelmingly negative (95% confidence intervals for the % contribution of persistent savings to overall: [−119% to −10%]; 40-trial washout data: [−92% to 6%]; 800-trial washout data [−135% to −12%]). The near- or complete absence of savings in temporally-persistent adaptation indicates that savings in the overall adaptation is primarily driven by temporally-volatile savings, and further suggests that temporally-volatile savings may be the sole source of savings in overall adaptation. Thus, our result indicates that savings arises from the faster reacquisition of volatile memories, rather than the re-manifestation of persistent memories.

### Temporally-volatile savings arise from implicit adaptation

Previous research associated savings in visuomotor adaptation with the rapid recall of explicit strategies, rather than faster implicit adaptation [18,30,31]. This led us to investigate the contributions of implicit and explicit processes in the temporally-volatile savings we observed in our paradigm. We thus ran Experiment 3 (N=40), which consisted of two 80-trial learning episodes separated by 800 washout trials. We dissected savings into implicit and explicit components using special instruction trials which prompted participants to disengage any explicit strategy by aiming their hand directly to the target [31,53–58].These instructions were presented immediately before and after the first (trial 10) 60s time delay following the onset of the visuomotor rotation in both initial learning and relearning, and allowed us to dissect adaptation into four subcomponents: implicit-persistent; implicit-volatile; explicit-persistent; explicit-volatile (Figure 4B, see Materials and Methods for details).

**Figure 4.**
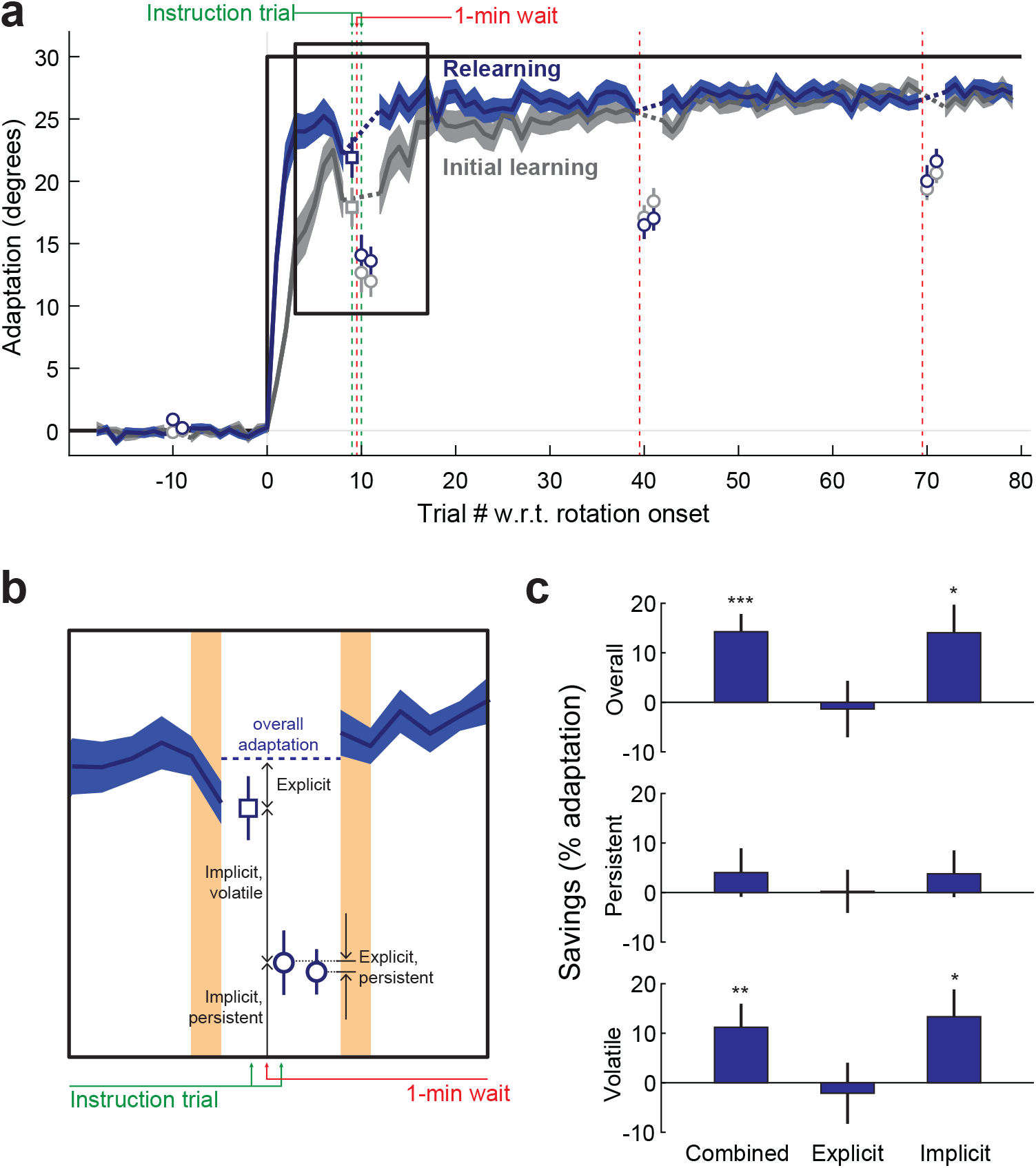
Temporally-volatile savings are due to an implicit adaptation component. **(a)** Learning curves for Experiment 3, showing overall adaptation (solid lines), overall implicit adaptation (square) and persistent adaptation (circles). Gray denotes initial learning, blue denotes relearning. Shading indicates SEM. **(b)** Detail from panel **a** illustrating the measurements and calculations involved in estimating overall, overall implicit, implicit-persistent, implicit-volatile and explicit-persistent components. Instructions one trial before and one after the 1-minute wait, allow the direct measurement of overall implicit and implicit-persistent components, respectively, in turn allowing the calculation of implicit-volatile as the difference between the two. The following trial is a non-instruction trial, allowing the direct measurement of explicit-persistent adaptation. **(c)** Savings in overall, temporally-persistent, and temporally-volatile adaptation in Experiment 3, both for combined implicit and explicit adaptation (left column) and broken into implicit and explicit components. In line with Experiments 1 and 2, data show overall and volatile, but not persistent, savings for combined (implicit + explicit) adaptation. Further dissociation into implicit and explicit components reveals this savings is due to implicit, temporally-volatile adaptation. Errorbars indicate SEM. * p<0.05; ** p<0.01.

In line with our findings in Experiments 1 and 2, we found savings for overall and volatile adaptation (14.3±3.6%, t(39) = 4.0, p = 0.00014 and 11.2±4.8%, t(37) = 2.4, p = 0.0119, correspondingly) but not persistent adaptation (4.0±4.9%, t(38) = 0.8, p = 0.21). Dissection of savings into explicit and implicit components revealed savings for both overall implicit and implicit-volatile adaptation (14.1±5.7%, t(38) = 2.5, p = 0.0088 and 13.3±5.5%, t(37) = 2.4, p = 0.0104, correspondingly) but not explicit-volatile adaptation (−2.1±6.2%, t(37) = −0.3, p = 0.63) or any of the persistent subcomponents (implicit-persistent: 3.8±4.7%, t(38) = 0.8, p = 0.21; explicit-persistent: 0.2±4.4%, t(38) = 0.1, p = 0.48). This finding suggests that overall savings were driven by the implicit and temporally-volatile component of adaptation, in turn suggesting that the temporally-volatile savings we observed in Experiments 1 and 2 predominantly reflect an implicit process rather than an explicit strategy. That the volatile component observed in Experiments 1 and 2 is primarily implicit is not surprising: First, it is unclear why an explicit strategy could be temporally-volatile to the point of being largely or completely forgotten after a short 1-minute delay. In fact, our recent work indicates that explicit adaptation displays essentially no temporal volatility, with over 95% stability across 1-minute delays[59]. Second, our paradigm elicited scant explicit adaptation (likely due to elements of our experiment design aimed at inducing implicit learning such as the use of point-to-point (rather than shooting) movements, the lack of aiming instructions, the lack of markers that could aid off-target aiming, and the presence of low-latency online feedback [56,60–63]), and without substantial explicit adaptation we lacked power for measuring explicit savings.

### Dissecting long-term memory in visuomotor adaptation

We next investigated whether the ability to dissect motor learning into temporally-persistent and temporally-volatile components could shed light on the mechanisms for the formation of long-term memories. To accomplish this, we examined the relationship between the levels of temporally-persistent and temporally-volatile learning observed after initial training and the amount of retention observed 24 hours later (Experiment 3). After a baseline period, we trained 25 participants on a 30° VMR for 120 trials. After this initial training, they were tested for temporally-persistent adaptation, retrained for 60 trials, and retested before returning the following day when they were tested for retention (**Fig. 5a**, see Materials and Methods).

**Figure 5.**
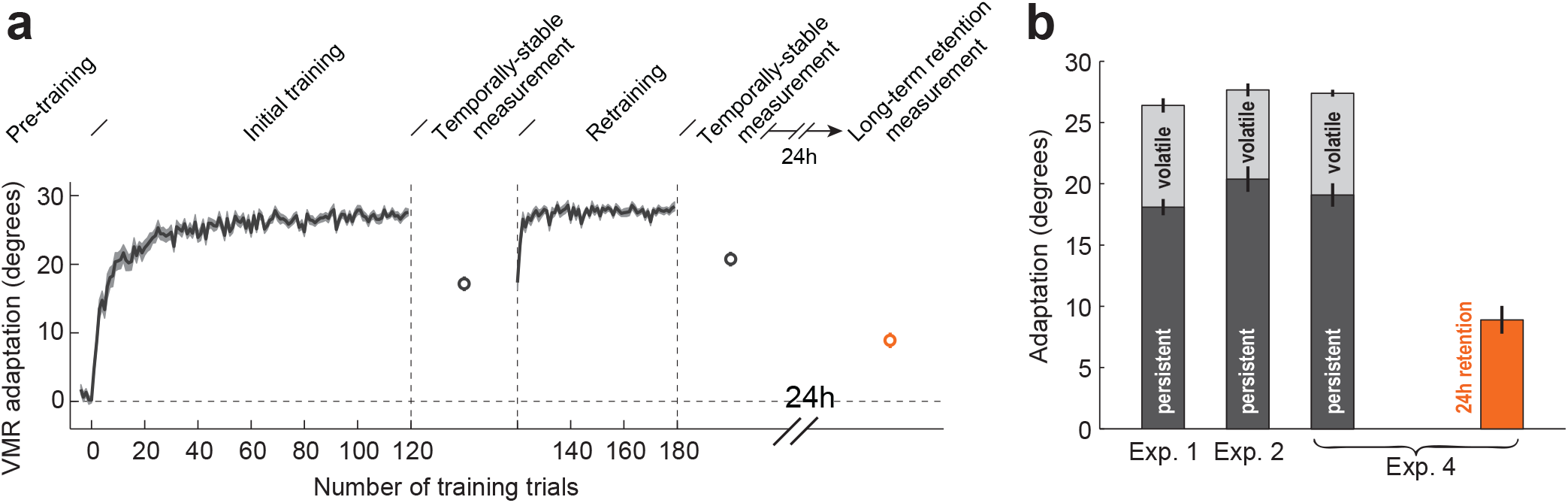
Temporally-persistent adaptation leads to long-term memory. **(a)** Experiment schedule and raw data for Experiment 4. After a baseline period, subjects were trained with a 30° VMR for 120 trials, and were then tested for temporally-persistent adaptation, retrained for 60 trials, and then retested. Subjects returned the following day when they were tested for 24-hour retention (orange circle). Note that 24-hour retention is lower than overall or temporally-persistent adaptation but higher than zero. **(b)** Comparison of overall, temporally-persistent, and temporally-volatile adaptation from Experiments 1, 2, and 4 with the 24-hour retention from Experiment 4. Experiments 1, 2, and 4 display similar levels of persistent and volatile adaptation.

We found that, the overall adaptation measured late in training in Experiment 3 (see Materials and Methods) was similar to that observed in Experiments 1 and 2 (27.4±0.3° for Experiment 3 vs. 26.4±0.6° and 27.7±0.5° for Experiments 1 and 2, see **Fig. 5b**). Similarly, the persistent component of adaptation was also similar across the three experiments (19.1±1.0° for Experiment 3 vs. 18.1±0.7° and 20.4±1.0° for Experiments 1 and 2, see **Fig. 5b**), suggesting that the somewhat longer training duration in Experiment 3 had little effect on either overall or temporally-persistent adaptation. When examining long-term memory, retained 24 hours after training, we found that participants retained 9.1±1.2° of the trained 30° rotation (orange bar in **Fig. 5b**). This corresponded to 32.4±4.2% of the overall learning and 45.6±4.6% of the temporally-persistent learning from day 1.

### Dissociable effects of temporally-volatile and temporally-persistent adaptation on the formation of long-term memory

To examine whether the dissection of day 1 learning into temporally-persistent and temporally-volatile components could shed light on the mechanism for long-term motor memory formation, we compared the levels of temporally-volatile, temporally-persistent, and overall learning to the amount of 24-hour retention for each individual participant. Looking for positive contributions of each component to 24-hour retention (using linear regression with regression coefficients restricted to be positive), we found no significant relationship between overall learning on day 1 and 24-hour retention on day 2 (r = +0.14, F(23,1) = 0.4, p = 0.51). However, we found a highly significant positive relationship between persistent learning on day 1 and 24-hour retention (slope = 0.80, r = +0.71, F(23,1) = 22.9, p = 0.00008). In contrast, we found no positive relationship between volatile learning on day 1 and 24-hour retention; in fact, the best-fit slope was zero (r=0.0, F(23,1) = 0, p=1), as the best fit slope without restricting regression coefficients would have been negative. This indicates that temporally-volatile learning does not lead to 24-hour retention, consistent with the fact that volatile learning, by definition, will over the course of one minute. We thus find that, whereas neither overall adaptation nor the temporally-volatile component can predict it, the temporally-persistent component of adaptation, measured only one minute after training, is able to accurately predict retention 24 hours after training.

We next performed a stepwise bivariate regression analysis of how 24 hour retention depended on temporally-volatile and temporally-persistent learning from day 1, as illustrated in **Fig. 6a,b**. This analysis was particularly important here because temporally-volatile and temporally-persistent learning were not independent across individuals but instead displayed a strong negative relationship such that participants with higher day 1 temporally-volatile learning displayed smaller day 1 temporally-persistent learning and vice versa. This bivariate regression revealed that adding temporally-volatile learning as a second regressor after temporally-persistent learning resulted in no significant improvement in the ability to explain 24-hour retention (R^2^ increased from 49.8% to 51.7% corresponding to a partial R^2^ of only 3.8%, F(22,1) = 0.4, p = 0.36). In contrast, adding temporally-persistent learning as a second regressor after temporally-volatile learning resulted in a large improvement in the ability to explain 24-hour retention (R^2^ increased from 0.0% to 51.7%, corresponding to a partial R^2^ of 51.7%, F(22,1) = 23.6, p = 0.00007). The results of this analysis are shown in **Fig. 6a,b** where we illustrate the partial R^2^ analysis by comparing each component of day 1 learning with the portion of 24-hour retention not explained by the other (See Materials and Methods for details).

**Figure 6.**
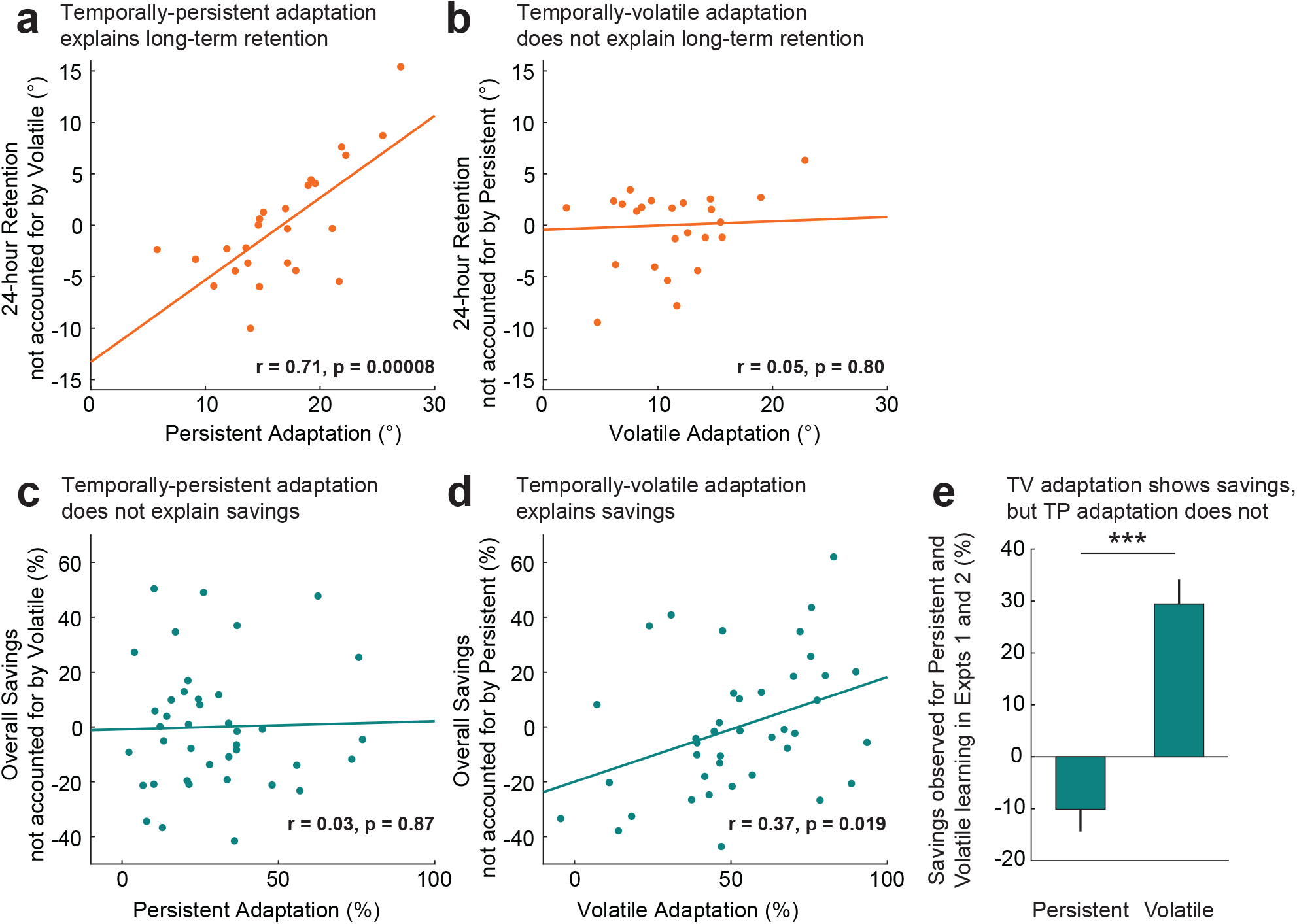
A Double dissociation between savings and long-term memory, uncovered by dissection of learning into temporally-persistent and temporally-volatile components. **(a)** Illustration of the partial regression between temporally-persistent adaptation on day 1 (shown on x-axis) and 24-hour retention across individuals (N=25). The y-axis represents residuals of the univariate regression of 24-hour retention upon temporally-volatile adaptation. The positive relationship indicates that higher temporally-persistent adaptation explains higher long-term memory. The solid line indicates linear fit. **(b)** Same as **a** but for temporally-volatile adaptation, showing no significant relationship (slopes restricted to positive values). **(c)** Illustration of the partial regression between temporally-persistent adaptation during relearning (shown in x-axis) and savings for early (trials 2-6) training combined across Experiments 1 and 2. The y-axis represents residuals of the univariate regression of savings upon temporally-volatile adaptation. There is no significant relationship. **(d)** Same as c but for temporally-volatile adaptation, showing a significant positive relationship and thus indicating that temporally-volatile adaptation explains overall savings. Note that mean y-axis values in **(a)-(d)** are zero, since they represent residuals of linear regression; these values do not reflect the actual amounts of long-term retention or savings which are, on average, significantly above zero. **(e)** Comparison of average savings for temporally-persistent (TP) and temporally-volatile (TV) adaptation, combined across Experiments 1 and 2. Errorbars indicate SEM. ***p<0.001.

In summary, we find in Experiment 4, that increased temporally-persistent adaptation leads to stronger long-term memory, whereas increased temporally-volatile adaptation does not. This sharply contrasts with Experiments 1-2 where we found that temporally-volatile adaptation led to savings whereas temporally-persistent adaptation did not. Taken together, these results demonstrate a striking double dissociation between the contributions of temporally-persistent and temporally-volatile learning to long-term memory and savings. To even more directly compare the two contributions to this double dissociation, we returned to the savings data from Experiments 1-2 and performed a bivariate analysis of the inter-individual differences in overall savings based on the levels of temporally-volatile and temporally-persistent learning during retraining (See Materials and Methods for details). This bivariate regression analysis is analogous to that performed on the 24-hour retention data above and is illustrated in **Fig. 6c,d**, in parallel format to **Fig. 6a,b**. Adding temporally-volatile learning as a second regressor after temporally-persistent learning resulted in a significant improvement in the ability to explain savings (R^2^ increased from 0.0% to 13.8% corresponding to a partial R^2^ of 13.8%, F(37,1) = 5.9, p = 0.0197). We note here that this p-value, corresponds to the variance reduction associated with adding temporally-volatile learning to the regression and is equivalent to a two-tailed test for its regression slope being non-zero. Thus, if a one-tailed test would have instead been used, in line with the idea of testing for a significantly positive relationship between savings and initial temporally-volatile learning, its p-value would have been < 0.01. In contrast, adding temporally-persistent learning as a second regressor after temporally-volatile learning resulted in no significant improvement in the ability to explain savings (R^2^ increased from 13.7% to 13.8%, corresponding to a partial R^2^ of 0.2%, F(37,1) = 0.1, p = 0.81). The findings from this regression analysis show that interindividual differences in the temporally-volatile but not the temporally-persistent component of initial learning explain individual differences in the amount of savings during relearning. This adds to the evidence illustrated in **Fig. 3**, summarized in **Fig. 6e**, that temporally-volatile learning displays savings whereas temporally-persistent learning does not.

## Discussion

Here we compared the mechanisms responsible for savings and long-term memory in human motor learning, finding that temporally-volatile adaptation leads to savings and that temporally-persistent adaptation leads to long-term memory. When we dissected adaptive responses into temporally-persistent and temporally-volatile components using 60-second breaks (**Fig. 1**), we found that temporally-persistent memories washed out 4-20x more slowly than temporally-volatile memories (**Fig. 2**), leaving a considerable temporally-persistent residual even after 100 washout trials (see **Fig. 2**), when overall adaptation had long since washed out to zero. This suggests that the short washout periods of 11-100 trials per trained movement direction used in a number of previous studies to wash out overall adaptation[13,15,16,18,23,48] would likely have failed to wash out this temporally-persistent adaptation.

We, therefore, controlled for the effect of residual temporally-persistent adaptation, either experimentally with an extended 800-trial washout period which could eliminate it, or analytically with appropriate baseline subtraction and normalization, allowed us to accurately assess savings. With this control, we consistently found significantly greater savings for temporally-volatile than temporally-persistent adaptation (**Figs. 3, 6**). In fact, whereas temporally-volatile adaptation showed savings by displaying relearning that was consistently faster than initial learning, temporally-persistent adaptation remarkably showed an anti-savings - displaying relearning that was significantly *slower* than initial learning both in the overall data and specifically in the 800-trial washout condition in which complete washout of both persistent and volatile learning occurred, allowing savings to be most cleanly measured. Temporally-persistent learning was also nominally slower, though not significantly so, in the 40-trial washout data. Remarkably, we found that savings in temporally-volatile adaptation was sufficiently large to overcome the anti-savings in persistent adaptation, and still confer robust savings on overall adaptation. Moreover, we found that the temporally-volatile savings we observed were due to implicit rather that explicit learning (**Fig. 4**). Our data thus suggest that savings in overall adaptation is derived from implicit, temporally-volatile adaptation, representing an increased propensity to more rapidly form a temporally-volatile memory, rather than the reemergence of a temporally-persistent memory.

When we dissected adaptation into volatile and persistent components to examine the mechanisms for long-term memory, we found a strong positive relationship between 24-hour retention and temporally-persistent but not temporally-volatile adaptation (**Figs. 5, 6**). Together, our findings for savings and 24-hour retention delineate a powerful double dissociation whereby temporally-persistent learning leads to long-term memory but not savings, and temporally-volatile learning leads to savings but not long-term memory.

### Juxtaposition between savings in temporally-volatile and temporally-persistent adaptation explains differences in savings across previous studies

Our findings provide a resolution to the apparent discrepancy between previous studies which isolated implicit savings in adaptation, yet found either anti-savings[32] or savings[20]. The paradigm used in the former study likely promoted temporally-persistent implicit adaptation, which here we find to display anti-savings, because compared to the current findings which employed only one movement direction, the multiplicity of movement directions used would dramatically increase the temporal spacing between same-direction movements, limiting temporally-volatile adaptation – though the extent of this effect is unfortunately difficult to definitively evaluate because the inter-trial time intervals were not explicitly reported in the study. In contrast, the quickly-paced paradigm used in the latter study would permit greater accumulation of temporally-volatile adaptation, which here we find to display enough savings to overcome anti-savings in temporally-persistent adaptation. Other factors may also have affected the balance between temporally-volatile and temporally-persistent components to drive the savings vs anti-savings observed in these two studies.

This effect whereby temporally-volatile savings would be reduced when a larger number of target directions are present in the experiment design, because same-direction inter-trial time intervals would increase and force temporally-volatile adaptation to decay to a greater extent, would also predict reduced savings even in studies that did not isolate implicit adaptation. This prediction is indeed borne out in previous work, with studies using 4-8-target paradigms finding either less pronounced savings[13,19] or no savings at all[23], whereas studies using 1-target or 2-target paradigms[15,17,18] demonstrating considerable savings. Given the 16s time constant observed for the decay of temporally-volatile adaptation during VMR learning[44], the fairly rapid 3s inter-trial interval in our study would allow near-complete (83%) carryover of temporally-volatile adaptation from one trial to the next, whereas the 8-fold increase in inter-trial intervals expected for an 8-target study with similar trial pacing would allow only 23% carryover. This would dramatically reduce the effect of the temporally-volatile savings.

### Incomplete washout can contaminate the assessment of savings

Our experiments revealed that temporally-persistent adaptation requires a surprisingly long period to wash out – well above 100 trials and much longer than overall adaptation that combines volatile and persistent components and is effectively washed out in just 20-40 trials (**Fig. 2**). This occurs because a negative temporally-volatile adaptation acts to mask the enduring temporally-persistent component after 20-40 washout trials. Therefore, if the washout of temporally-persistent adaptation is not specifically measured, it is easy to get the false impression that a short block of 20-40 trials is sufficient to washout all adaptation so that true savings – which refers to relearning after complete washout of all adaptation – can be cleanly measured. In fact, most savings studies we examined employed washout periods well below what would be necessary for complete washout of temporally-persistent adaptation[13,15,16,18,23,48] – a notable exception being a recent [32] study which employed a long washout period with several breaks that probed for residual temporally-persistent adaptation. Incomplete washout of temporally-persistent adaptation would lead to an *apparent* savings due to the unmasking of this persistent memory during relearning as predicted by a multi-rate learning model[10], contaminating the assessment of *true* savings. Even Zarahn et al. [15], where the primary claim was that savings was present after the complete washout of all components of adaptation so that the savings they found could not be explained by *apparent* savings due to the unmasking of a slowly decaying learning process, used only 40 trials meaning that the savings they observed almost certainly included *apparent* savings from incomplete washout of temporally-persistent adaptation. The current results from the 800-trial washout data where temporally-persistent adaptation was entirely eliminated, do show clear evidence for the *true* savings that Zarahn et al hypothesized, and it is unfortunately impossible to know just how much of the savings they reported was due to incomplete washout because of paradigmatic differences that may have increased or decreased the degree to which the washout was incomplete.

### Implicit savings does not result from the recall of a consolidated motor memory

Our findings dissociating savings from long-term memory, and demonstrating savings to be driven by temporally-volatile, rather than temporally-persistent memories, upend the widespread view that the faster relearning that characterizes savings results from the recall of a previously consolidated, stable motor memory[8,9,14,16,17,19,30,64,65]. Whereas savings has been taken as a litmus test for the consolidation of motor memory 1, 2 or even 7 days after training[13,23], here we find that savings is primarily driven by the temporally-volatile component of adaptation, which decays over the course of one minute or less. These temporally-volatile savings could not arise from a consolidated memory of previous adaptation, as any memory solid enough to survive the hour-long, 800-trial washout period in our experiments, would certainly be stable against the passage of time during the minute-long breaks in our experiment used to define temporal lability. Our findings thus indicate that savings is not driven by a consolidated memory of the adapted motor output, in line with the double dissociation between savings and long-term memory that we demonstrate.

In contrast, the savings we observe is driven by an increased learning rate for adaptive but temporally-volatile changes in motor output from one trial to the next when experiencing the same perturbation again after washout. This ability is very different from the usual conception of a consolidated motor memory in which the trained actions themselves are remembered. Thus, our findings are consistent with a model in which savings is based on an ability to improve the adaptation of actions, rather than a memory for the actions themselves.

### Mechanisms for learning rate modulation in temporally-volatile adaptation

What are the mechanisms behind learning rate increases in temporally-volatile adaptation? Recent work suggests that such a learning rate increase can be driven by learning environments with increased statistical consistency, defined as a positive correlation between successive perturbations (i.e. lag-1 autocorrelation) in environmental dynamics or errors from one trial to the next[39–41]. This consistency-driven effect is further enhanced when the environment repeats the same perturbation[41], with highly consistent, repetitious switching environments increasing learning rates up to 3x from baseline and highly inconsistent ones decreasing then up to 5x. Critically, the initial training periods in savings paradigms, including the current one, are characterized by both consistency and repetition, as they usually consist of a large number of trials with the same perturbation, leading to an increase in the learning rate during retraining.

### Savings and explicit adaptation

Experiment 3 revealed that the temporally-volatile savings we observe arises from implicit adaptation. This adds to recent evidence [19,20] against the idea that savings is exclusively driven by explicit adaptation [18,30–32]. Although savings can clearly occur because of explicit strategies, we did not observe this in our experiment. While this may, in part, reflect the considerable variability in the balance between implicit and explicit adaptation across individuals [54,66], the lack of explicit savings in our study was likely due to an experimental design that promoted implicit adaptation and minimized explicit strategy, and thus provided little power to detect explicit savings. The design elements included the lack of aiming instructions, the absence of workspace markers that could aid re-aiming, the use of point-to-point rather than shooting movements, and the engineering of low (~25ms) visual feedback latency for onscreen cursor motion, all of which may promote implicit learning [56,60–63]. In contrast, studies which reversed most of these design elements elicited primarily explicit adaptation and found clear explicit savings [31].

### Parallels between temporally-volatile / temporally-persistent learning and the fast/slow learning processes of motor adaptation

Another line of work dissected motor adaptation, not experimentally, but instead on the basis of a computational model with two distinct adaptive processes: a fast adaptive process that learns rapidly and displays weak retention, and a slow adaptive process that learns slowly and displays strong retention[10]. By manipulating the training duration in order to elicit different amounts of the fast and slow learning, a subsequent study found that 24-hour retention was specifically predicted by the amount of slow learning, rather by the amount of fast learning or overall adaptation[67]. Interestingly, this model-based dissection mirrors our temporal-stability based dissection: the slow process, like temporally-persistent adaptation, leads to 24-hour retention, whereas the fast process, like temporally-volatile adaptation, does not. We note that in the Joiner et al. study, 49±6% (95% confidence) of slow learning on day 1 is retained after 24 hours, mirroring our Experiment 3 finding that 46±9% (95% confidence) of persistent learning on day 1 is retained after 24 hours. Moreover, the trial-to-trial learning characteristics of the fast and slow processes mirror the ones for volatile and persistent adaptation, respectively. In particular, slow adaptation displays slower learning and better retention than fast adaptation, just as temporally-persistent adaptation displays slower learning and better retention than temporally-volatile adaptation (see **Fig. 3c,d** and **Fig. 2a**, respectively). These parallels challenge the idea that the fast process corresponds to explicit adaptation [68], as they argue that the fast process corresponds to temporally-volatile adaptation which here we find to be implicit. Taken together these observations suggest that temporally-volatile adaptation is captured by the fast process of the two-state model, and temporally-persistent adaptation is captured by the slow process.

## Materials and Methods

A total of 106 subjects (44 men, age 22.2±4.1, 13 left-handed) participated in the present study (20 each in Experiments 1 and 2, 41 in Experiment 3, and 25 in Experiment 4). Participants were naïve with respect to the purpose of the experiments and provided written informed consent in accordance with the policies of the Harvard University Institutional Review Board.

### Apparatus

We used the same experimental setup as the one used in recent work[46,47]. Subjects sat in front of an apparatus consisting of a 200Hz digitizing tablet (Wacom Intuos 3 12”x19”, resolution of position data: 0.005mm; accuracy: 0.25mm) positioned below a 23” 120Hz LCD monitor. During the experiment, subjects moved a custom-made handle, which contained a stylus, on top of the tablet allowing us to record hand position. Vision of the hand was occluded by the monitor and subjects instead observed their movement on the screen through a white cursor representing hand position.

### Experiment protocol

Using their dominant hand, subjects made point-to-point arm reaching movements between a starting position and targets 9cm away. At the end of each movement, they were rewarded with a bell sound if they had managed to reach and stop at the target within 250ms. Training was isolated to the outward movements, as visual feedback was unavailable during the return movements. Subjects took rest breaks roughly every 200 trials (about 7-10 minutes, see **Fig. 1**).

Experiments 1 and 2 consisted of reaches towards a 90° target direction (in the midline, directly away from the body). After a 220-trial baseline block with no visual rotation, subjects in Experiment 1 (N=20) entered the main part of the session which contained three 80-trial training periods. During training, a 30° visuomotor rotation (VMR) was imposed about the starting position. The sign of this VMR was the same for all training periods for each subject, with half the subjects training with a clockwise VMR and the other half training with a counter-clockwise VMR. The first and second training periods were separated by a 40-trial washout period, whereas the second and third training periods were separated by an 800-trial washout period. The training schedule in Experiment 2 (N=20) was the same apart from that the 800-trial washout period came first (between the first and second training periods, see **Fig. 1**).

Experiment 3 (N=41) was similar to Experiment 2 in that it contained two 80-trial training periods separated by a 800-trial washout period (but not a third training period). It was designed to examine whether the temporally-volatile savings like the ones observed in Experiments 1 and 2 were due to an implicit or explicit process.

To dissect savings into implicit and explicit components, we used special instruction trials which prompted participants to disengage any explicit strategy by aiming their hand directly to the target. This method, also referred to as *exclusion* (since participants are to exclude strategies from their reach)[69], has been, in various forms, widely used to dissect implicit and explicit visuomotor adaptation[31,53–58]. Specifically, instructions were given to either move to the center of the target or to its near/far end (both of which would not alter the reaching angle) and were presented immediately before and after the first (trial 10) 60s time delay within both visuomotor rotation training episodes (initial learning and relearning).

This enabled us to directly assess overall implicit adaptation (the amount of adaptation on the first instruction trial) and implicit-persistent adaptation (the amount of adaptation in the second instruction trial, which followed a 60s delay); and, by comparing these two, this enabled us to assess implicit-volatile adaptation. Moreover, by comparing adaptation in the second instruction trial to the no-instruction trial following it, we assessed explicit-persistent adaptation; and, by estimating overall adaptation as the average adaptation two trials before and after all these delay/instruction trials, we obtained estimates of overall explicit, volatile, and persistent adaptation (Figure 4B).

To minimize delays in reaction time, which would increase the inter-trial time interval and lead to further reduction in temporally-volatile adaptation, participants were presented with an “upcoming instruction” sound during the trial preceding the instruction. To familiarize participants with instruction trials (and the preceding “upcoming instruction” sound) ahead of visuomotor rotation training, we presented a series of similar instruction trials during familiarization. Familiarization contained four different possible instructions: move your hand to the near, far, left or right end of the (circular) target. There were clear biases towards the instructed endpoints showing adherence to the instructions.

The aim of Experiment 4 (N=25) was to examine the formation of long-term memories of VMR adaptation. The experiment began with a baseline period with no VMR that consisted of 456 trials, spread evenly across 19 target directions. After this baseline, subjects were trained on a 30° VMR for 120 reaches to a target placed at 90° (in the midline, directly away from the body, see **Fig. 4a**). The direction of the 30° visual rotation was approximately balanced, with 13 subjects trained with a counter-clockwise VMR and 12 subjects with a clockwise VMR. This was followed by a testing block with 3 reaches towards each of the 19 targets. During this block, visual feedback was withheld so that repeated measurements could be made without these measurements being contaminated by additional training that could be elicited by visual feedback. We used the first movement towards the training direction to measure temporally-persistent adaptation. After this testing block, subjects were retrained on the 30° VMR for an additional 60 trials and after that were tested again without visual feedback to measure temporally-persistent adaptation as described above. Participants returned the following day to be tested for 24-hour retention without visual feedback.

### Sample size determination

While sample sizes for experiment groups in analogous studies typically range between 8 and 12, here we used somewhat larger sample sizes (N = 20, 20, 41 and 25 for Experiments 1, 2, 3 and 4, respectively). For Experiments 1 and 2, we examined a larger number of participants so that we could rigorously assess not only whether savings is present or not for temporally-persistent and temporally-volatile adaptation, but also the time-course of savings for these two adaptation components at multiple points during training, as well as whether there are any subtle differences in savings or the extent of washout following the 40-trial vs the 800-trial washout periods. The larger sample sizes in Experiments 1 and 2 also enabled more precise comparisons between the time course of washout for temporally-persistent and temporally-volatile adaptation, as the time constant estimates for these washout curves can be especially susceptible to noise in the data. In Experiment 3, we doubled the sample size relative to Experiment 2, given that Experiment 3 involved dissection of adaptation into four (explicit-persistent, explicit-volatile, implicit-persistent, implicit-volatile), rather than two components. In Experiment 4, we examined N=25 participants as we wanted to be able to look at not just the group-average amount of 24-hour retention, but also examine how inter-individual differences in 24-hour retention on day 2 related to inter-individual differences in temporally-persistent and temporally-volatile adaptation on day 1 (**Fig. 4b,c**).

### Data analysis

#### Statistical comparisons

We performed single-sided paired t-tests across subjects to assess the presence of (positive) savings in adaptation and its subcomponents. For all other statistical comparisons two-sided paired t-tests across subjects were implemented, with the exception of the comparisons involving the estimation of washout time constants in **Fig. 2a** and the estimation of confidence intervals associated with the % contribution of temporally-persistent or temporally-volatile savings to overall savings: in these cases we used a bootstrapping procedure (see below) instead of comparing fits to individual subject data, because the high noise in these individual data leads to low confidence about the corresponding individual parameters.

#### Data inclusion criteria

We performed outlier rejection on the learning curves of each experiment. Specifically, for each trial, we excluded adaptation levels that were more than 3 inter-quartile ranges (IQRs) away from the subject median. This resulted in the inclusion of 99.4% of trials.

#### Estimation of visuomotor rotation adaptation

To assess the amount of adaptation to the trained VMR, we measured the direction of hand motion on each trial. In movements with visual feedback, this was defined as the direction of the vector between the hand position at movement onset (based on a 6.4 cm/s velocity threshold), and the hand position 150ms later. We used 150ms to measure feedforward adaptation, as feedback corrections should be minimal at this point. In movements with no visual feedback, this was defined as the direction of the vector between the hand position at movement onset and the movement endpoint. To examine learning-related changes in performance, we subtracted out the small bias present in the baseline (0.13±0.11°), from all the movement-direction data.

#### Measurement of temporally-persistent and temporally-volatile adaptation

In Experiments 1, 2 and 3, we measured temporally-persistent adaptation using one-minute breaks interspersed with training. Because the temporally-volatile component of motor adaptation decays with a time constant of 15-20 seconds[42,44], the one-minute breaks we impose here amount to 3-4τ, and thus lead to a 95-98% decay in temporally-volatile adaptation, effectively isolating the temporally-persistent component of adaptation. In contrast, the trial-to-trial decay in temporally-volatile adaptation for non-break trials would be much lower, as the experiments were fast-paced with a median inter-trial time interval of 2.5-2.7s, amounting to 0.1-0.2τ thus leading to only 10-15% decay. We thus used the amount of adaptation on the trial immediately following such a break as a measure of the temporally-persistent adaptation on that trial. These timed one-minute breaks occurred every 30 trials during the VMR training blocks (on trials 10, 40, and 70 after the onset of each 80-trial training episode), and in 40-trial intervals during the long washout period, as shown in **Fig. 1b,c**. During these breaks subjects held the handle still on the starting position.

In addition to these timed one-minute breaks, Experiments 1 and 2 contained a number of additional rest breaks which allowed subjects to put the handle aside and were not strictly timed. These breaks occurred only during baseline or washout periods as shown in **Fig. 1c**. We used the amount of adaptation after these breaks as a measure of temporally-persistent adaptation on the corresponding trials as above, but only in cases where these breaks resulted in inter-trial intervals greater than 40s (65.8% of these breaks).

Finally, we measured volatile adaptation for the same trials by first estimating the corresponding amount of overall adaptation around each persistent adaptation measurement, calculated as the average adaptation two trials before and two trials after the post-break trial on which persistent adaptation was measured. We then calculated temporally-volatile adaptation as the difference between these overall and temporally-persistent adaptation measurements.

In Experiment 4, temporally-persistent adaptation was measured during the no-feedback testing blocks that followed rest breaks (average break duration: 124±9s, minimum 48s) so that temporally-persistent adaptation could be assessed using the three reaches in each block that were towards the training target. We estimated volatile adaptation as the difference between persistent adaptation and overall adaptation. The latter was assessed as the average adaptation during the last 20 trials of the training and retraining blocks. Finally, we calculated 24-hour retention based on the no-feedback data from the testing block on day 2 (**Fig. 5a**).

#### Estimation of washout time constants

The washout of overall adaptation proceeded in two timescales: a very rapid initial washout phase during the first 2-3 washout trials, during which adaptation levels went from about 27° to about 11°, and then a slower washout phase that is illustrated in **Fig. 1c**. To compare the time constants for washout for both temporally-persistent and overall adaptation (**Fig. 2a**), we focused our overall washout analysis on the period beginning at trial 3 of washout in order to focus on the slower washout phase for comparing overall and temporally-persistent washout, because no temporally-persistent measurements were available during the very fast initial phase.

To estimate the values and confidence intervals associated with the time constants for washout, τ, we utilized a bootstrapping procedure[70]. Specifically, for each one of 10000 bootstrap iterations, we randomly sampled, with replacement, N=20 subjects from each group, and fit their average data with a single-exponential fit (Equation 1):

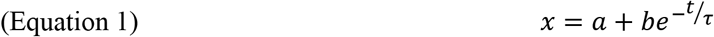

When analyzing the overall washout curves we discarded not only trials after each one-minute or rest break that removed the temporally-volatile component of adaptation in order to measure temporally-persistent learning, but also the three trials immediately thereafter, during which temporally-volatile adaptation might not be fully reequilibrated.

#### Normalization of adaptation data

To systematically quantify savings, and specifically take into account the systematically different baselines between post-long washout vs. post-short washout relearning, as well as the different baselines between temporally-persistent, temporally-volatile and overall adaptation, we subtracted baseline adaptation, *X_baseline_*, and normalized each learning curve *x* by the distance between baseline and the ideal adaptation level of 30 degrees (Equation 2). The baseline level for overall adaptation was defined as the average of the last 5 trials before training onset, whereas the baseline level for persistent adaptation was defined as the average of the last three persistent-adaptation trials before training onset (in the case of baselines for initial training and training after an 800-trial washout) or as the last single persistent-adaptation trial, trial, 10 trials before the onset of training (in the case of baselines for training after a 40-trial washout, since the 40-trial washout contained only a single persistent-adaptation measurement trial).

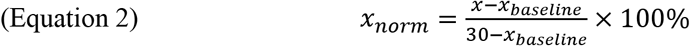

#### Estimation of savings

Finally, savings for each type of adaptation were taken to be the % difference in adaptation between initial training and retraining for the same training trial (Equation 3).

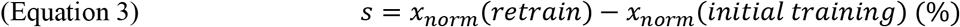

Throughout the study, we focused on savings around 1-minute delay trials (especially trial 10 after training onset which captured early adaptation, but also trials 40 and 70), as these were the trials for which all three types of adaptation could be assessed. For the analysis of the across-individual relationships between savings and persistent/volatile adaptation (**Fig. 6c,d**), however, because the measurement of temporally-volatile adaptation was based on the same measurements as overall adaptation (volatile = overall [adaptation two trials before and after the 60s wait trial] – persistent [adaptation on the 60s wait trial]), we instead calculated overall savings based on trials 2-6 in order to ensure that any observed relationships were not due to measurements shared between the dependent (savings) and independent (temporally-volatile adaptation) variables. This range was selected as it was both relatively far from the measurements used to calculate temporally-volatile adaptation, but also better captured the rapid rise of overall adaptation providing more power to assess inter-individual differences in savings.

#### Comparisons of inter-individual differences

To examine contributions of temporally-persistent or temporally-volatile adaptation on savings and long-term memory (**Fig. 6a-d**), we used linear regression with slopes restricted to positive values to model positive contributions of these components of adaptation and either savings or long-term retention. Specifically, for studying long-term memory, we compared temporally-persistent and temporally-labile adaptation on day 1 in Experiment 4 against 24-hour retention on day 2, whereas, for studying savings, we compared temporally-persistent and temporally-labile adaptation from trial 10 in the retraining blocks in Experiments 1 and 2 against overall savings calculated as in the preceding paragraph.

## Acknowledgements

We would like to thank Jasmine Bailey and Emerson Fang for their help with the experiments; Jordan Brayanov, Joel Greenwood and Edward Soucy for their help with instrumentation; Rene Chen for the illustration in Figure 1a; Andrew Brennan and Yohsuke Miyamoto for helpful discussions.

## Author contributions

AMH and MAS designed experiments, analyzed the data and wrote the paper. JRM designed experiments and edited the paper.

